# Quantitative variations of ADF/cofilin’s multiple actions on actin filaments with pH

**DOI:** 10.1101/422824

**Authors:** Hugo Wioland, Antoine Jegou, Guillaume Romet-Lemonne

## Abstract

Actin Depolymerizing Factor (ADF)/cofilin is the main protein family promoting the disassembly of actin filaments, which is essential for numerous cellular functions. ADF/cofilin proteins disassemble actin filaments through different reactions, as they bind to their sides, sever them, and promote the depolymerization of the resulting ADF/cofilin-saturated filaments. Moreover, the efficiency of ADF/cofilin is known to be very sensitive to pH. ADF/cofilin thus illustrates two challenges in actin biochemistry: separating the different regulatory actions of a single protein, and characterizing them as a function of specific biochemical conditions. Here, we investigate the different reactions of ADF/cofilin on actin filaments, over four different values of pH ranging from pH 6.6 to pH 7.8, using single filament microfluidics techniques. We show that lowering pH reduces the effective filament severing rate by increasing the rate at which filaments become saturated by ADF/cofilin, thereby reducing the number of ADF/cofilin domain boundaries, where severing can occur. The severing rate per domain boundary, however, remains unchanged at different pH values. The ADF/cofilin-decorated filaments (refered to as “cofilactin” filaments) depolymerize from both ends. We show here that, at physiological pH (pH 7.0 to 7.4), the pointed end depolymerization of cofilactin filaments is barely faster than that of bare filaments. In contrast, cofilactin barbed ends undergo an “unstoppable” depolymerization (depolymerizing for minutes despite the presence of free actin monomers and capping protein in solution), throughout our range of pH. We thus show that, at physiological pH, the main contribution of ADF/cofilin to filament depolymerization is at the barbed end.

A number of key cellular processes rely on the proper assembly and disassembly of actin filament networks ^1^. The central regulator of actin disassembly is the ADF/cofilin protein family ^2, 3^, which comprises three isoforms in mammals: cofilin-1 (cof1, found in nearly all cell types), cofilin-2 (cof2, found primarily in muscles) and Actin Depolymerization Factor (ADF, found mostly in neurons and epithelial cells). We refer to them collectively as “ADF/cofilin”.

Over the years, the combined efforts of several labs have led to the following understanding of actin filament disassembly by ADF/cofilin. Molecules of ADF/cofilin bind stoechiometrically ^4, 5^ to the sides of actin filaments, with a strong preference for ADP-actin subunits ^6^–^10^. Though ADF/cofilin molecules do not contact each other ^11^, they bind in a cooperative manner, leading to the formation of ADF/cofilin domains on the filaments ^5, 7, 9, 12, 13^. Compared to bare F-actin, the filament portions decorated by ADF/cofilin (refered to as “cofilactin”) are more flexible ^14, 15^ and exhibit a shorter right-handed helical pitch, with a different subunit conformation ^11, 16^–^19^. Thermal fluctuations are then enough to sever actin filaments at (or near) domain boundaries^8, 9, 13, 20, 21^. Cofilactin filaments do not sever, but depolymerize from both ends ^13^ thereby renewing the actin monomer pool.

ADF/cofilin thus disassembles actin filaments through the combination of different actions. As such, it vividly illustrates a current challenge in actin biochemistry: identifying and quantifying the multiple reactions involving a single protein. This is a very difficult task for bulk solution assays, where a large number of reactions take place simultaneously, and single-filament techniques have played a key role in deciphering ADF/cofilin’s actions ^9, 13, 20, 22^–^24^. In particular, the microfluidics-based method that we have developed over the past years, is a powerful tool for such investigations ^25^. It has recently allowed us to quantify the kinetics of the aforementioned reactions, and to discover that ADF/cofilin-saturated filament (cofilactin) barbed ends can hardly stop depolymerizing, even when ATP-G-actin and capping protein are present in solution ^13^.

In addition, ADF/cofilin is very sensitive to pH ^4, 5, 26^–^29^. In cells, pH can be a key regulatory factor ^30^. It can vary between compartments, between cell types, and be specifically modulated. We can consider that a typical cytoplasmic pH would be comprised between 7.0 and 7.4. Recently, we have quantified the different reactions involving ADF/cofilin at pH 7.8 ^13^, leaving open the question of how these reaction rates are indivdually affected by pH variations. For instance, it has been reported that ADF/cofilin is a more potent filament disassembler at higher pH values ^4, 5, 26^–^29^ but the actual impact of pH on the rate constants of individual reactions has yet to be characterized. Moreover, whether the unstoppable barbed end depolymerization that we have recently discovered for ADF/cofilin-saturated filaments at pH 7.8 ^13^ remains significant at lower, more physiological pH values is an open question.

Here, we investigate how the different contributions of ADF/cofilin (using unlabeled ADF, unlabeled cof1 and eGFP-cof1) to actin filament disassembly depend on pH, which we varied from 6.6 to 7.8. We first present the methods which we have used to do so, based on the observation of individual filaments, using microfluidics (Fig. 1). We measured cofilin’s abitility to decorate actin filament by binding to its sides (Fig. 2), and the rate at which individual cofilin domains severed actin filaments (Fig. 3). We next quantified the kinetic parameters of filament ends, for bare and ADF/cofilin-saturated (cofilactin) filaments (Fig. 4), and we specifically quantified the extent to which the barbed ends of cofilactin filaments are in a state which can hardly stop depolymerizing (Fig. 5). We finally summarize our results (Fig. 6).

## METHODS

### Buffers and proteins

Experiments were carried out at room temperature in F-buffer (10 mM Hepes or Tris-HCl, 50 mM KCl, 1 mM MgCl_2_, 0.2 mM EGTA, 0.2 mM ATP, 10 mM DTT and 1 mM DABCO) with different pH values: pH 6.6 (Hepes), pH 7.0 (Hepes or Tris), pH 7.4 (Tris), and pH 7.8 (Tris). The pH of each buffer was adjusted after mixing all the ingredients.

All the protocols for protein purification can be found in reference ^13^. Actin was purified from acetone poweder, made from rabbit muscle. Recombinant mouse cofilin-1, eGFP-cof1 (with eGFP at the N-terminus), human ADF, human profilin-1, and human gelsolin were expressed in bacteria and purified. Capping protein was made from recombiant mouse capping proteins alpha1 and beta2. Spectrin-actin seeds were purified from human erythrocytes.

Gelsolin was biotinylated with sulfo-NHS-biotin. Actin was fluorescently labeled on accessible surface lysines of F-actin, using Alexa 488 or Alexa-568 succimidyl ester.

### Microfluidics for the study of single actin filaments

In order to distinguish the different actions of cofilin, and quantify them, one needs to observe single events on individual actin filaments. To do so, the microfluidics-based method that we have developed over the past 9 years ^25^ is a valuable tool. The microfluidics setup is sketched in Fig. 1A. It allows one to monitor a large number of actin filaments anchored by one end only, as well as labeled cofilin, using epifluorescence or TIRF microscopy (Fig. 1B). The experiments we report here are very similar to the ones we performed in reference ^13^. Barbed end dynamics, as well as cofilin side-binding and filament severing, were monitored on filaments grown from spectrin-actin seeds anchored to the coverslip surface (Fig. 1B). Pointed end dynamics were monitored by anchoring gelsolin-capped barbed ends do the coverslip surface, using biotinylated gelsolin and a neutravidin-decorated surface.

**Figure 1.**
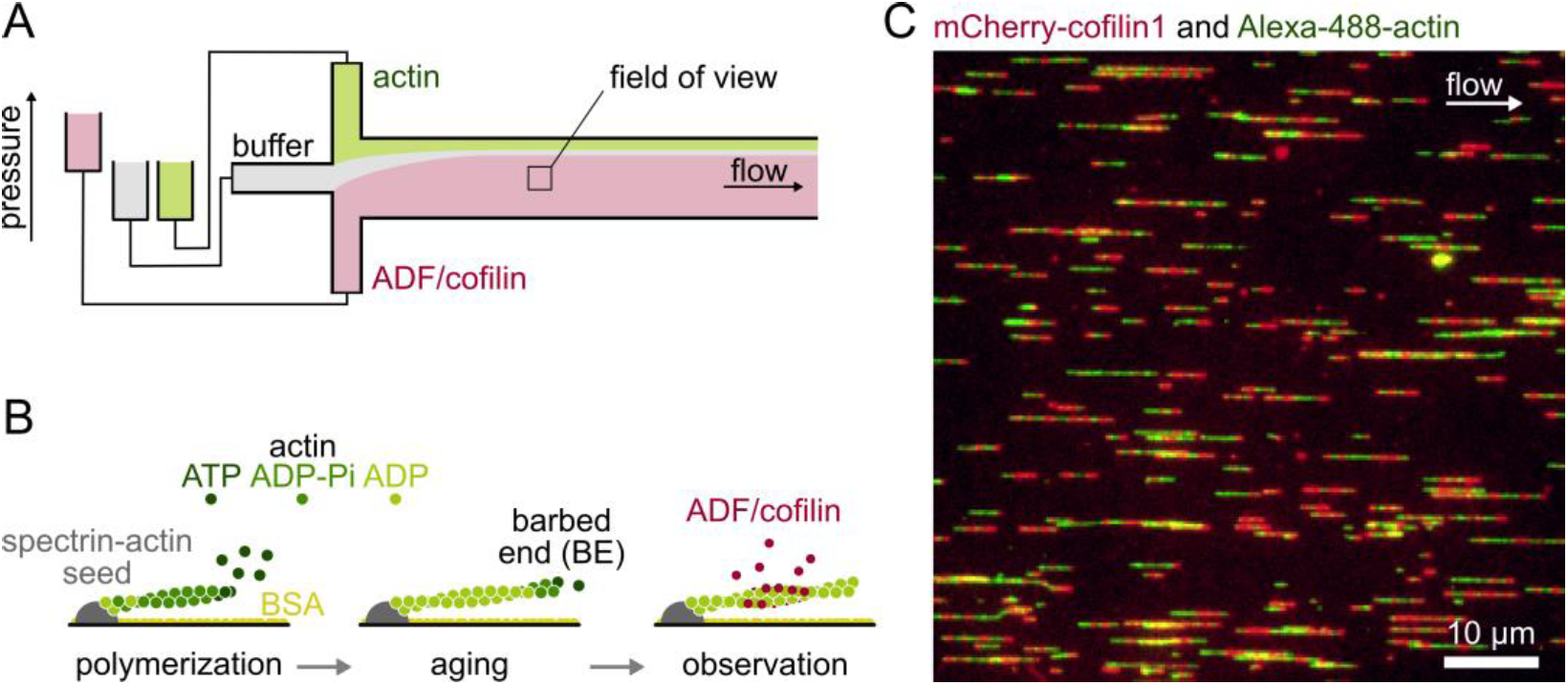
Using microfluidics to monitor individual actin filaments and the binding of cofilin. (A) Experiments are performed in microfluidic chambers, sketched from above. The main channel is connected through three inlets to different protein solutions. Controlling the pressure in each inlet allows one to rapidly change the solution in the field of view. (B) Sketch of a typical experiment (side view). Filaments are elongated from coverslip-anchored spectrin-actin seeds, by flowing in with ATP-G-actin. Filaments are then aged by flowing in a solution of ATP-G-actin at the critical concentration, for at least 15 min. This results in >99% of the monomers in the ADP-state. Finally, filaments are exposed to ADF/cofilin. (C) Example of a field of view, imaged with TIRFm. ADP-F-actin labelled with Alexa-488 is exposed to mCherry-cofilin-1, which forms observable domains on the filaments.

### Microscopy and data analysis

Images were acquired in epifluorescence or TIRF microscopy (ILAS2, Roper Scientific, now Gataca systems) on a Nikon TiE inverted microscope equiped with a 60x oil-immersion objective, either with an Evolve EMCCD camera (Photometrics) controlled by Metamorph, or with an Orca-Flash2.8 camera (Hamamatsu) controlled by micromanager. Images were analyzed using ImageJ.

Elongation or depolymerization rates (Fig. 4) were determined on individual filaments, and median values were reported. We considered that each actin subunit contributed 2.7 nm to the filament length. For the quantification of severing (Fig. 3), uncapping (Fig. 5C) and rescue (Fig. 5E, F), survival fractions were determined, following a Kaplan-Meier algorithm^31^.

**Figure 2.**
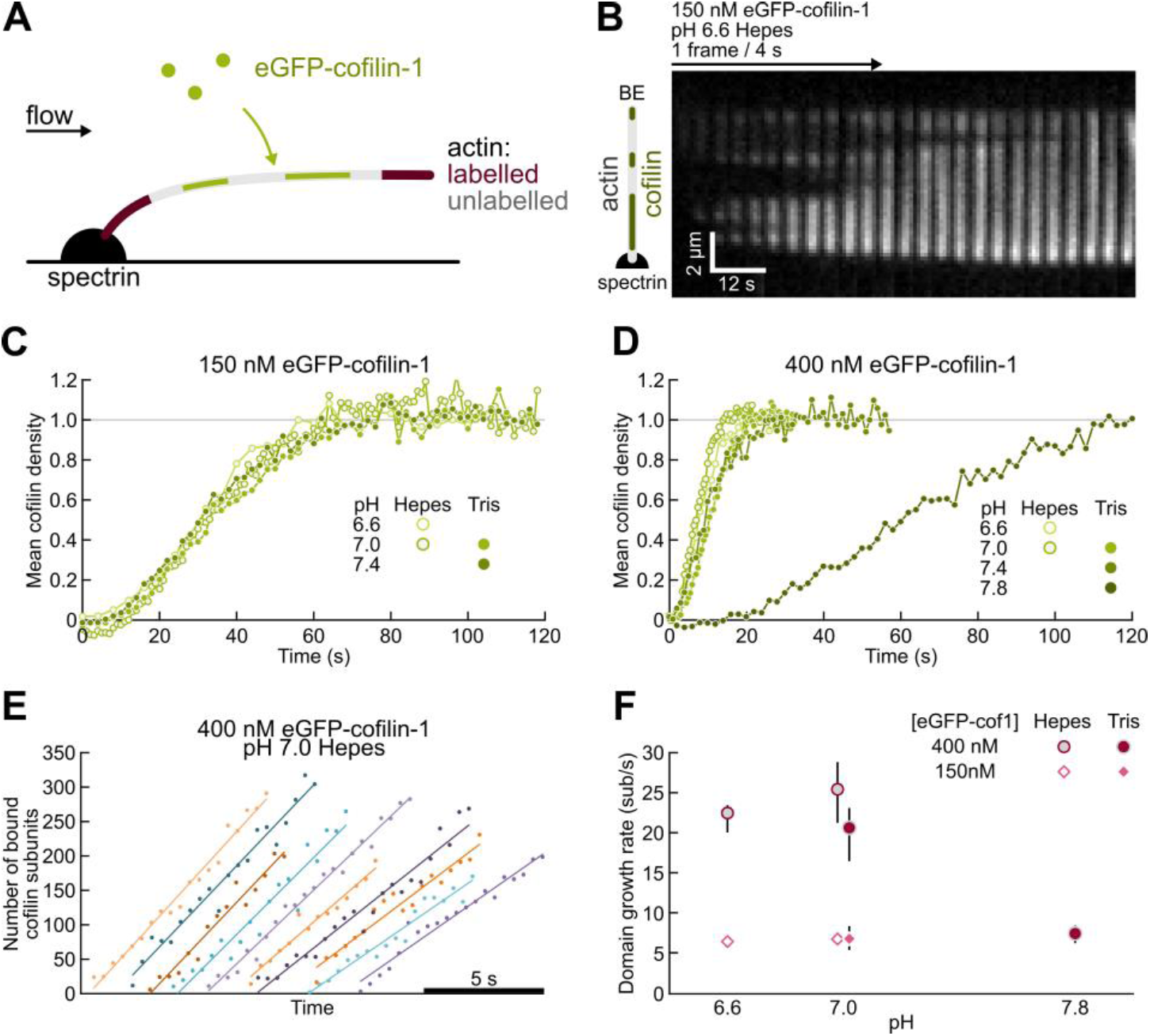
Cofilin binds more slowly to filaments at higher pH values. (A). Experimental configuration. Actin filaments are grown from spectrin-actin seeds with a long middle segment of unlabelled ADP-actin. (B). Time-lapse showing an unlabeled ADP-actin filament become saturated by eGFP-cof1 over time. (C-D). Mean normalized eGFP-cofilin-1 fluorescence signal, binding onto unlabelled ADP-F-actin. 150 nM (C) and 400 nM (D) eGFP-cofilin-1 was injected in the chamber from time t=0 onwards. The fluorescence signal was averaged along 20 to 35 pixels (3.5 to 6 µm) for each filment. Number of filaments (C) N = 10, 10, 18, 20, for pH 6.6 Hepes, 7.0 Hepes, 7.0 Tris and 7.4 Tris, respectively, and (D) N = 10 in all conditions. (E). Number of cofilin subunits in individual domains, increasing over time. For clarity, the time origin has been shifted for each curve. Lines: linear fit. Condition: 400 nM eGFP-cofilin-1, pH 7.0 Hepes. (F). Growth rate of individual cofilin domains at different eGFP-cofilin-1 concentrations and pH. Value: median, error bars: interquartile range. N = 10 domains, except N = 9 for pH 6.6 Hepes 150 nM cof1, and for pH 7.8 Tris 400 nM cof1.

**Figure 3.**
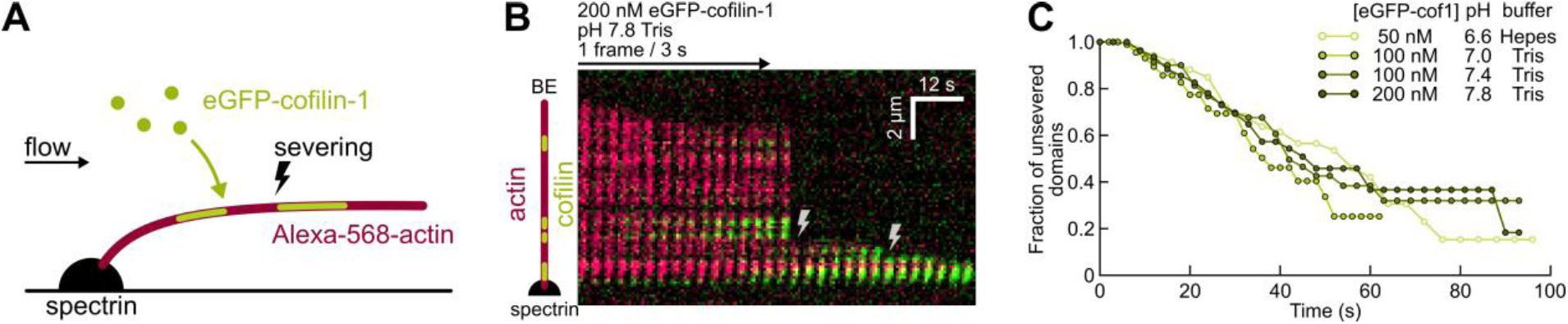
The severing rate per cofilin domain is unaffected by pH. (A). Experimental setup. Alexa-568-labelled actin filaments are polymerized from actin-spectrin seeds, and aged before being exposed to eGFP-cof1. (B). Typical kymograph. 200 nM eGFP-cof1 (green) is constantly injected, binds F-actin (red) and induces severing (lightning symbols). (C). Fraction of cofilin domains with no severing event detected near their edges, over time. Time t=0 is defined for each domain as the last frame before they become visible. The survival fraction curves are calculated using the Kaplan-Meier method over 22 to 43 filaments, 78 to 90 cofilin domains and 30 to 33 severing events, for each data set.

**Figure 4.**
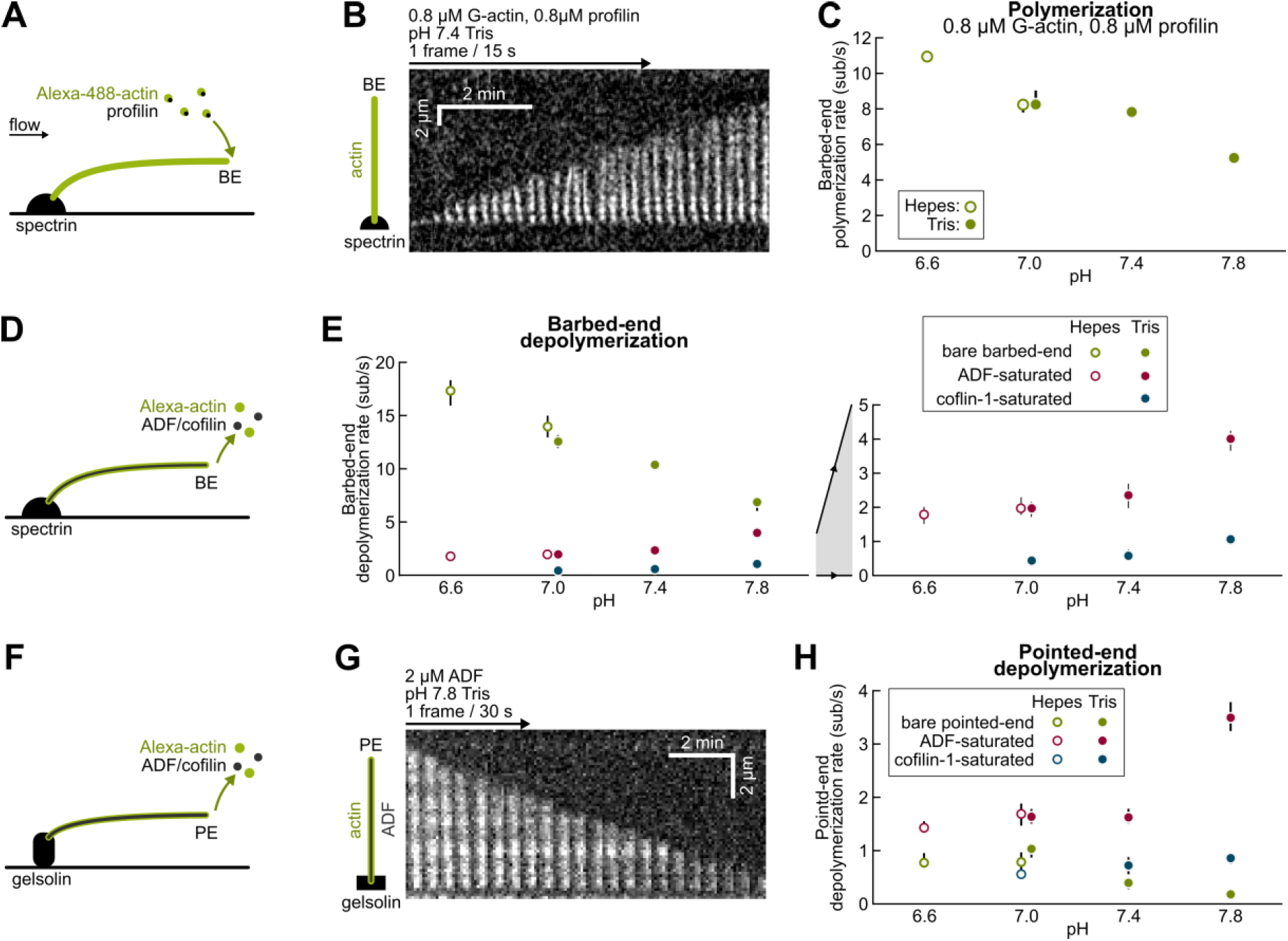
Higher pH slows down polymerization and depolymerization of bare F-actin but accelerates depolymerization of ADF/cofilin-saturated filaments at both ends. (A-C) Polymerization from the barbed-end. (A) Sketch of the experimental configuration, where filaments were grown from actin-spectrin seeds with G-ATP-actin and profilin. (B) Kymograph of a typical elongating filament. (C) Polymerization rate at different pH. N = 20 filaments for each condition. (D-E) Depolymerization from the barbed-end. (D) Sketch of the experimental configuration. ADP-F-actin is exposed either to buffer only, or to 1-2µM unlabelled ADF or cofilin-1 in order to fully saturate the filament in less than a minute. (E) Depolymerization rate for different pH values. Right: zoom into the 0-5 sub/s range. From left to right, N = 20, 32, 22, 32, 31 (buffer only); N=9, 14, 23, 33, 34 (ADF-saturated); N=17, 18, 16 (cofilin-1-saturated). (F-H) Depolymerization from the pointed-end. (F) Sketch of the experimental configuration. ADP-F-actin was bound to the surface by gelsolin. Filaments were exposed to buffer only (supplemented with 0.4 mM CaCl_2_ to ensure gelsolin-actin tight binding), containing 1 to 2 µM unlabelled ADF or cofilin-1 to rapidly saturate filaments. (G) Typical kymograph of a depolymerizing filament saturated with ADF. (H) Pointed-end depolymerization rate at different pH. N = 14, 20, 15, 20, 20 (buffer); N=20, 20, 16, 20, 20 (ADF-saturated); N= 20, 20, 20 (cofilin-1-saturated). (C, E, H) Symbol: median, error bars: interquartile range.

**Figure 5.**
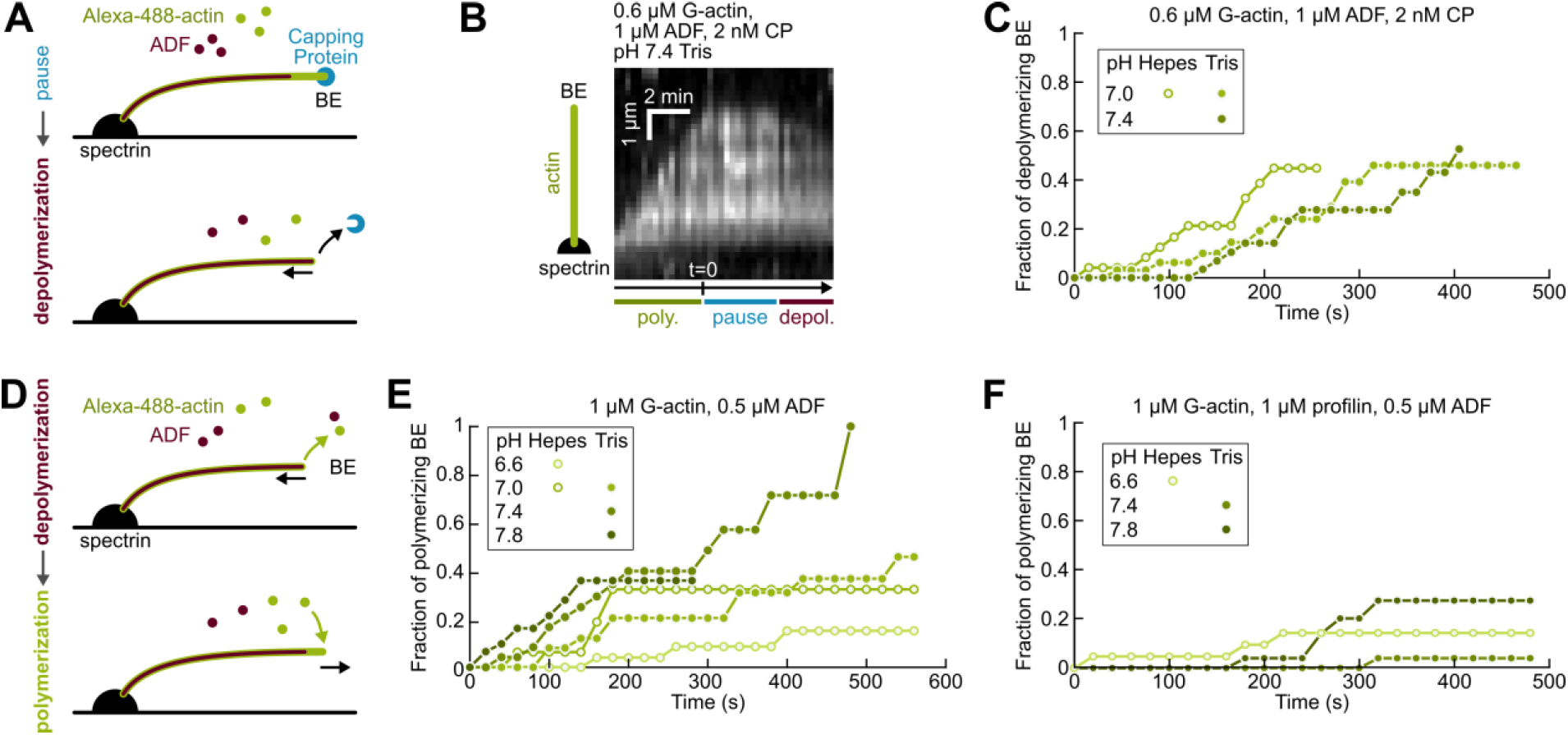
The “unstoppable” depolymerization of cofilactin barbed ends is observed throughout our pH range. (A-C) Synergy of CP and ADF/cofilin to saturate filaments and initiate barbed end depolymerization. (A) Sketch of experimental configuration and events. Filaments grow until they are capped with CP. ADF/cofilin can then saturate the filaments, up to their BE which thus uncaps and depolymerizes. (B) Kymograph of a filament continuously exposed to the same solution containing 0.8 µM G-ATP-actin, 1 µM ADF and 2 nM CP. The filament polymerizes, pauses as it is capped by CP, and eventually depolymerizes. (C) Fraction of barbed ends that transitioned from a pause to depolymerization. Time t = 0 corresponds to the beginning of the pause (as shown on B). N = 24, 32, 32 filaments for pH 7.0 Hepes, pH 7.0 Tris, pH 7.4 Tris, respectively. (D-F) Cofilactin barbed ends sustain depolymerization in the presence of ATP-G-actin. (D) Sketch of the experimental configuration and events. Filaments are polymerized from spectrin-actin seeds and saturated with ADF. Depolymerizing cofilactin filaments are then constantly exposed to a solution of ATP-G-actin. (E) Fraction of barbed ends that transitioned from depolymerization to polymerization over time, when exposed to 1 µM ATP-G-actin and 0.5 µM ADF (to keep filaments saturated). N = 25, 16, 25, 24, 31 for pH 6.6 Hepes, 7.0 Hepes, 7.0 Tris, 7.4 Tris, 7.8 Tris, respectively. (F) Same as (E), with 1 µM profilin added to the solution. N= 21, 27, 30 for pH 6.6 Hepes, 7.4 Tris, 7.8 Tris, respectively.

**Figure 6.**
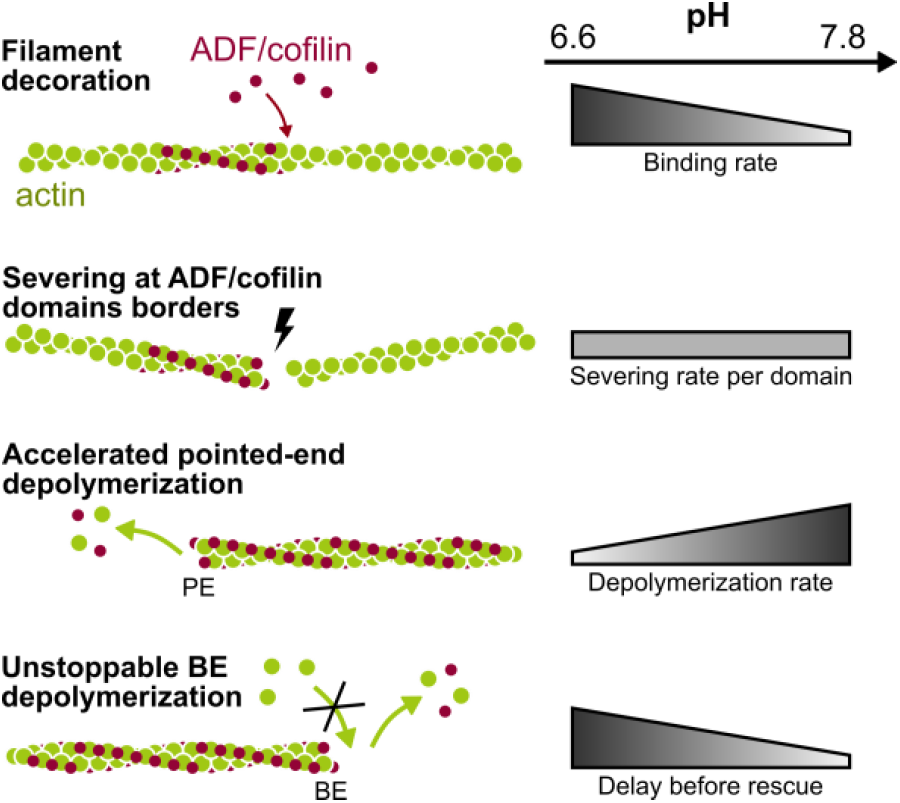
Summary of results: barbed end depolymerization is an important contribution of cofilin disassembly at physiological pH. Within the range of pH that we have explored (pH 6.6 to 7.8) we have made the following observations (from top to bottom, on this sketch). A lower pH favors the rapid decoration of filaments by ADF/cofilin, but the severing rate per cofilin domain does not vary with pH. As a consequence, at a higher pH, domain boundaries persist longer (before domains merge) and severing is more efficient. The acceleration of pointed end depolymerization for cofilactin filaments is mostly observed at high pH. The “unstoppable” depolymerization of cofilactin barbed ends is observed at all pH, and is more pronounced at lower pH values.

As an example, we detail here the protocol for the quantification of severing (Fig. 3). Filaments were polymerized from anchored spectrin-actin seeds with Alexa-568(14%)-actin and aged for at least 15 min to ensure that > 99% of the monomers were in a ADP-state ^25^. A solution of low concentration eGFP-cofilin-1 in F-buffer (no G-actin) was then constantly injected. Images were acquired using epifluorescence microscopy. All domains located at least 0.5µm (4 pixels) away from the anchored seed were analyzed. For each domain, time t=0 was defined as the frame before which they appeared. Domains could then ‘sever’, i.e. have a filament severing event occur near one of their boundaries, or be ‘lost’, for example when a severing event occured at another domain located upstream on the same filament. These events were accounted for using a Kaplan-Meier algorithm in order to determine the survival fraction of unsevered domains over time (Fig. 3C).

## RESULTS AND DISCUSSION

### Cofilin binds faster to actin filaments at lower pH values

Using our microfluidics setup we have generated ADP-actin filaments comprising a long unlabeled segment, and we have monitored the binding of eGFP-cof1 to this segment (Fig. 2A, B). We found that the decoration of the filament was equally fast at pH 6.6, 7.0 and 7.4 (Fig. 2C), but significantly slower at pH 7.8, where it took approximately 6 times longer to reach 50% of full saturation in the presence of 400 nM eGFP-cof1 (Fig. 2D).

We mesured the growth rate of individual eGFP-cof1 domains (Fig. 2E), and found that they appeared to grow symmetrically toward both filament ends, as we already reported for pH 7.8 ^13^. Similar to our observation for the overall decoration of filaments, we found that domain growth rate was pH-independent for low values of pH, and was approximately 3-fold lower at pH 7.8 (Fig. 2F). Overall, our results show that cofilin domains nucleate and grow much faster at pH 6.6-7.4 than at pH 7.8.

### The severing rate per cofilin domain is unaffected by pH

We next sought to measure the severing rate per eGFP-cof1 domains, at different pH values. To do so, we exposed Alexa-568 (14%) ADP-actin filaments to eGFP-cof1, and monitored the severing events over time, for each cof1 domain (Fig. 3). As previously reported, severing events were observed to occur at the boundaries of cof1 domains, and occurred more often at the pointed end side of the domain. Different cofilin concentrations were used at different pH values in order to observe separate, individual domains long enough (domains grow faster and thus fuse more rapidly at lower pH values). We found no significant differences in the severing rate per domain, as a function of pH (Fig. 3C).

Therefore, our results indicate that the previously reported enhancement of filament severing activity by cofilin at higher pH values ^32^ does not come from a faster severing at each potential severing site, but from a greater number of these sites, i.e. a greater number of domain boundaries. For a given concentration range, the rapid cofilin decoration at low pH makes domain boundaries less numerous and shorter-lived, as domains rapidly expand and merge.

### Bare actin filaments (without cofilin) are more dynamic at lower pH values

Before measuring the depolymerization rates of ADF/cofilin-saturated filaments, we measured the barbed end elongation rate as well as the depolymerization rate of both ends in the absence of ADF/cofilin, at different pH values (Fig 4). We found that barbed ends exhibited higher on- and off-rates at low pH (Fig. 4C, E) and that pointed ends also had higher off-rates at low pH (Fig. 4H). This is consistent with earlier work on pH ^33^–^35^, and studies perfomed at high pH values ^25^ typically report slower filament dynamics than studies performed at lower pH values ^36^.

### Cofilin-saturated filaments depolymerize faster at higher pH values

The pointed end depolymerization of ADF- and cof1-saturated filaments is faster at higher pH values (Fig. 4H). As a result, the enhancement of pointed end depolymerization by ADF-saturation, which is very significant at pH 7.8 (a 17-fold increase, compared to bare filaments) is milder at physiological pH (a 4-fold increase at pH 7.4 and a 2-fold increase at pH 7). Cof1-saturated filament pointed ends at physiological pH (7.0-7.4) depolymerize at rates similar to those of bare filaments. This effect likely contributes to the more efficient filament disassembly previously reported for higher pH values ^4, 5, 26 – 29^ (in addition to severing, which we have discussed earlier in this manuscript).

When filaments were saturated with cof1 or ADF, barbed ends depolymerized slower than those of bare filaments, and their off-rate increased with pH (Fig. 4E). Consequently, the difference in barbed end depolymerization between bare and saturated filaments was greater at lower pH values: ADF-saturated barbed ends depolymerized 6.6-fold slower than bare barbed ends at pH 7, but only 1.7-fold slower at pH 7.8.

Another, totally different effect of ADF/cofilin on barbed end dynamics, is that free ADF/cofilin molecules in solution directly target bare ADP-actin barbed ends and increase the monomer off-rate, as we have first reported at pH 7.8 ^13^. This effect remains true at lower pH (Supp. Fig. S1). This effect, which requires to have ADP-actin at the barbed end, is unlikely to play a role in cells, where an ATP-actin monomer will quickly bind the barbed end and thus protect it from the direct targeting by ADF/cofilin ^13^. Moreover, this enhancement of depolymerization by direct targeting of the barbed end disappears if the sides of the filament are decorated with ADF/cofilin up to the barbed end. At physiological pH 7-7.4, the faster decoration of the filament sides by ADF/cofilin (Fig. 2) make this direct targeting of the BE even less likely to play a role in cells.

Nonetheless, the effect of direct BE targeting by ADF/cofilin can be readily observed in vitro, if actin monomers are absent from solution and the barbed end thus remains ADP-actin (Supp Fig. S1). Importantly, this effect should not be confused with the saturation of the sides of the filaments with ADF/cofilin, which slows down barbed end depolymerization (Fig. 4E). This was unfortunately the case in a recent study ^37^ where the authors, using unlabeled ADF, wrongly concluded that binding ADF to the sides of filaments accelerated their depolymerization from the barbed end.

### The barbed ends of cofilactin filaments are even harder to stop depolymerizing at lower pH values

We next investigated if the unstoppable barbed end depolymerization of ADF/cofilin-saturated filaments, which we discovered at pH 7.8 ^13^, was also true at lower pH values.

We verified that, at physiological pH (7-7.4), capped filaments exposed to ADF became uncapped and started depolymerizing (Fig. 5A-C). Since lower pH values accelerate the formation and growth of cofilin domains on filaments (observed for cof1, Fig. 2) they also reduce the time required for these domains to reach the barbed end and uncap it: at pH 7.0 and 7.4 (Fig. 5C) uncapping occurs faster than what we previously observed at pH 7.8 ^13^. In order to further quantify the unstoppable nature of barbed end depolymerization for ADF/cofilin-saturated filaments, we compared the time it took for 1µM ATP-G-actin to “rescue” ADF-saturated filament barbed ends from depolymerization. We found that this rescue was slower at lower pH values (Fig 5E). We found that adding profilin in the buffer delayed further the rescue of depolymerizing barbed ends (Fig. 5F).

Our results thus show that the “unstoppable” depolymerizing state of cofilactin barbed ends is a feature that exists over the whole pH range that we have explored. In fact its contribution to the depolymerization of cofilactin filaments appears to be greater at physiological pH (7-7.4) than at pH7.8 where it was originally discovered ^13^ for the following reasons: (1) depolymerizing cofilactin barbed ends are more difficult to rescue at lower pH; (2) capped actin filaments are more rapidly saturated by ADF/cofilin and uncapped at lower pH; and (3) ADF-saturated filaments depolymerize faster from their barbed ends than their pointed ends, which depolymerize almost as slow as the pointed ends of bare actin filaments at lower pH.

## CONCLUSIONS

Our results further illustrate the power of single-filament microfluidics as a tool for actin biochemistry. We could not have obtained these results with bulk solution assays, and microfluidics offered a number of advantages compared to standard single filament techniques ^25, 38, 39^. Here, it allowed us to distinguish and to separately quantify the main reactions of ADF/cofilin on actin filaments, for different pH values: binding to the sides (Fig. 2) and severing (Fig. 3) of actin filaments, and, for filaments decorated by ADF/cofilin, the acceleration of pointed end depolymerization (Fig. 4) and the unstoppable depolymerization of barbed ends (Fig. 5). The quantitative variations of these reactions with pH are summarized in Figure 6.

Overall, our results are consistent with the notion that lowering the pH mostly affects the conformation of the actin filament, which is then more favorable for binding ADF/cofilin ^40^. Indeed, at lower pH we find that bare actin filaments are less cohesive and depolymerize faster, and that decoration by ADF/cofilin makes them more stable thanks to the additional bonds it provides. Our observation that ADF/cofilin binds more readily to actin filaments at lower pH values is also consistent with the idea that actin filaments are in a more “cofilin-friendly” conformation. These changes in F-actin conformation with pH likely involve the binding of cations to specific sites on the subunits, which modulate filament properties ^41^ and whose reorganization is coupled to cofilin binding ^42^.

Faster ADF/cofilin-binding at lower pH values also explains why previous studies have reported a weaker severing activity at lower pH values: domain boundaries, where severing can occur, rapidly vanish as domains rapidly expand and merge. We found that the severing rate per domain was unaffected by pH, within the range we explored. Importantly, we found that cofilactin filament pointed ends did not depolymerize much faster than bare filaments barbed end at physiological pH values. In contrast, the unstoppable barbed end depolymerization of cofilin-saturated filaments remains an important feature at all pH, and is even stronger for lower pH values. Our results thus show that, at physiological pH, the dominant effect differentiating cofilactin filaments from bare filaments is the nature of their barbed ends, and their “unstoppable” depolymerization.

## AUTHOR INFORMATION

Corresponding authors: Antoine Jegou and Guillaume Romet-Lemonne.

Funding: We acknowledge funding from the Human Frontier Science Program (grant RGY0066 to G.R.-L.), French Agence Nationale de la Recherche (grant Muscactin to G.R.-L.), European Research Council (grant StG-679116 to A.J.), Fondation ARC pour la Recherche sur le Cancer (postdoctoral fellowship to H.W.).

## ACKNOWLEDGEMENTS

We thank all members of the Romet/Jegou lab, and especially Bérengère Guichard and Sandy Jouet for help with buffers and protein purification.

## ABBREVIATIONS

ADF, Actin Depolymerizing Factor; BE, barbed end; PE, pointed end; F-actin, filamentous actin; G-actin, globular (monomeric) actin; cofilactin, cofilin(or ADF)-decorated actin filament; TIRFm, Total Internal Reflection Fluorescence microscopy.

**Supplementary Figure S1.**
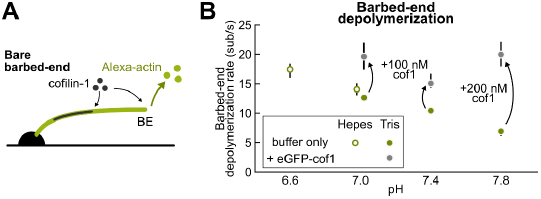
Depolymerization of bare barbed ends is accelerated by ADF/cofilin in solution. (A)Sketch of the experimental configuration. ADP-F-actin is exposed either to buffer only or to eGFP-cofilin-1 at low concentration. Labelled cofilin-1 was used to ensure that the cofilin domains on the filament did not reach the BE. (B)Depolymerization rate for different pH values. From left to right, N = 20, 32, 22, 32, 31 (buffer only); N=10 for all (+eGFP-cofilin-1). The data points with buffer only (green) are the same as in Fig.4E.

